# Direct Comparison of Lysine vs. Site-specific Protein Surface Immobilization in Single-molecule Mechanical Assays

**DOI:** 10.1101/2023.03.17.532846

**Authors:** Haipei Liu, Zhaowei Liu, Mariana Sá Santos, Michael A. Nash

## Abstract

Single-molecule force spectroscopy (SMFS) is a powerful method for studying folding states and mechanical properties of proteins, however, it requires surface immobilization of proteins onto force-transducing probes such as cantilevers or microscale beads. A common immobilization method relies on coupling surface-exposed lysine residues to carboxylated surfaces using 1-ethyl-3-(3-dimethyl-aminopropyl) carbodiimide and N-hydroxysuccinimide (EDC/NHS). Because proteins typically contain many lysine groups, this strategy results in a heterogeneous distribution of tether positions in the molecule. Genetically encoded peptide tags (e.g., ybbR) provide alternative chemistries for achieving site-specific immobilization, but thus far a direct comparison of site-specific vs. lysine-based immobilization strategies to assess effects on the observed mechanical properties was lacking. Here, we directly compared lysine- vs. ybbR-based protein immobilization in SMFS assays using several model polyprotein systems. Our results show that lysine-based immobilization results in significant signal deterioration for monomeric streptavidin-biotin interactions, and loss of the ability to correctly classify unfolding pathways in a multipathway Cohesin-Dockerin system. We developed a mixed immobilization approach where a site-specifically tethered ligand was used to probe surface-bound proteins immobilized through lysine groups, and found partial recovery of specific signals. The mixed immobilization approach represents a viable alternative for mechanical assays on *in vivo*-derived samples or other proteins of interest where genetically encoded tags are not feasible.

## Introduction

Single-molecule force spectroscopy (SMFS) with the atomic force microscope (AFM) can characterize dissociation pathways of receptor-ligand complexes under force, resolve intermediate unfolding states in globular domains, and quantify energy landscapes governing conformational transitions in proteins. To implement this technique, a protein of interest is stretched using a force transducing probe, typically a magnetic or optically-trapped microscale bead or nanofabricated cantilever tip. Due to the vectorial nature of force, the positions within the molecule that are used for tethering the molecule of interest to the surface strongly influence the mechanical response^[1–6]^, therefore the bioconjugation method used for protein immobilization is critically important for interpreting the resulting force response. Early work relied on non-specific adsorption of multi-domain polyproteins containing independently foldable domains in a single polypeptide chain, and was used to probe N-to-C pulling geometries^[7–11]^. When probing receptor-ligand interactions, however, non-specific adsorption as an immobilization technique is typically not suitable because it does not allow sufficient differentiation between specific and non-specific signals and because pick-up of ligand molecules can clog receptors on the tip.

To address this issue, methods relying on covalent bond formation to immobilize proteins for receptor-ligand SMFS were developed^[12]^. For example, poly(ethylene) glycol (PEG) polymeric chains were used to provide a flexible linker to attach molecules onto the AFM tip. PEG can minimize non-specific adsorption and present the molecules tens to hundreds of nanometers away from the surface to preserve binding activity. A common approach uses PEG-based linkers to immobilize proteins by coupling to primary amine groups on lysine residues in the native sequence ^[13–16]^. This approach of randomly linking lysine groups to PEG is convenient because it is compatible with commercially available proteins and polymers, however, one limitation is that it results in a diversity of pulling orientations on the molecule of interest.

Alternatively, several site-specific protein ligation methods relying on genetically encoded peptide sequences or ligatable domains have been reported for immobilization of biomolecules for force spectroscopy. Examples include the SNAP Tag^[17,18]^, HaloTag^[19–22]^, Sortase^[23–25]^, SpyTag/SpyCatcher^[26–29]^, histidine-tag/Ni^2+^-NTA^[30,31]^, maleimide-cysteine^[32]^, azide–alkyne cycloaddition^[6,33,34]^, asparaginyl endopeptidases^[35,36]^, and Sfp phosphopantetheinyl transferases in combination with the ybbR tag^[37–39]^. These site-specific protein immobilization techniques require custom protein synthesis to incorporate the tag, however, they enable precisely defined pulling points on the molecule of interest which can significantly improve the homogeneity of surface functionalization and specificity of measured signals. Meanwhile, an improved flexible linker of elastin-like polypeptides (ELPs)^[40]^ could be fused between the ligation site and the protein of interest, providing a uniform stretching response. Despite anecdotal reports of the improvements that can be achieved using site-specific linkage, thus far no direct comparison study exists. The aim of this work therefore was to directly compare random lysine coupling with site-specific immobilization in a set single-molecule mechanical assays on model polyproteins. We quantified the influence of site-specific immobilization for various molecular systems by directly comparing lysine to Sfp/ybbR tag coupling strategies and evaluating the impact on experimental outcomes in terms of precision, yield, and reproducibility.

### Lysine vs. site-specific linkage for monomeric streptavidin-biotin interactions

We first conducted AFM-SMFS on complexes consisting of a monomeric variant of streptavidin (mSA)^[41–43]^ bound to biotin, where mSA was immobilized on a glass surface and PEG-biotin was attached to a cantilever. As one of the first receptor-ligand systems measured in AFM-SMFS^[44]^, the streptavidin-biotin interaction has been well characterized while the newly developed monomeric variants with unambiguous binding stoichiometry shows slightly different rupture forces compared to those in tetramer form^[45,46]^. We attached identical fusion proteins of ybbR-ddFLN4-mSA to a glass surface through poly(ethylene) glycol (PEG) linkers using either EDC/Sulfo-NHS to couple to lysine residues in the fusion protein. Alternatively, we used Coenzyme A (CoA) together with Sfp phosphopantetheinyl transferase to couple to the N-terminal ybbR tag comprising the sequence DSLEFIASKLA. The fusion domain ddFLN4 refers to the 4th domain of the F-actin cross-linking filamin rod of Dictyostelium discoideum, a commonly used fingerprint domain in SMFS assays. The respective chemical schemes for lysine-based and site-specific protein immobilization onto the glass surface are shown in **Fig. 1**. For measurements on both lysine-based and site-specific immobilized ybbR-ddFLN4-mSA, biotin was covalently coupled to the cantilever through a PEG linker attached at the C5 carboxylic acid away from the tetrahydrothiophene ring in biotin. The mSA fusion protein studied here (**Fig. 2a**) contains 9 total lysine residues (6 in ddFLN4, 3 in mSA), each of which can serve as potential PEG linkage sites in the lysine-based method. The rupture force data for the lysine-based method therefore reflects an average of the mechanical responses of the system when anchored from lysines, however, we expect not all of the lysine residues are equally reactive.

**Figure 1.**
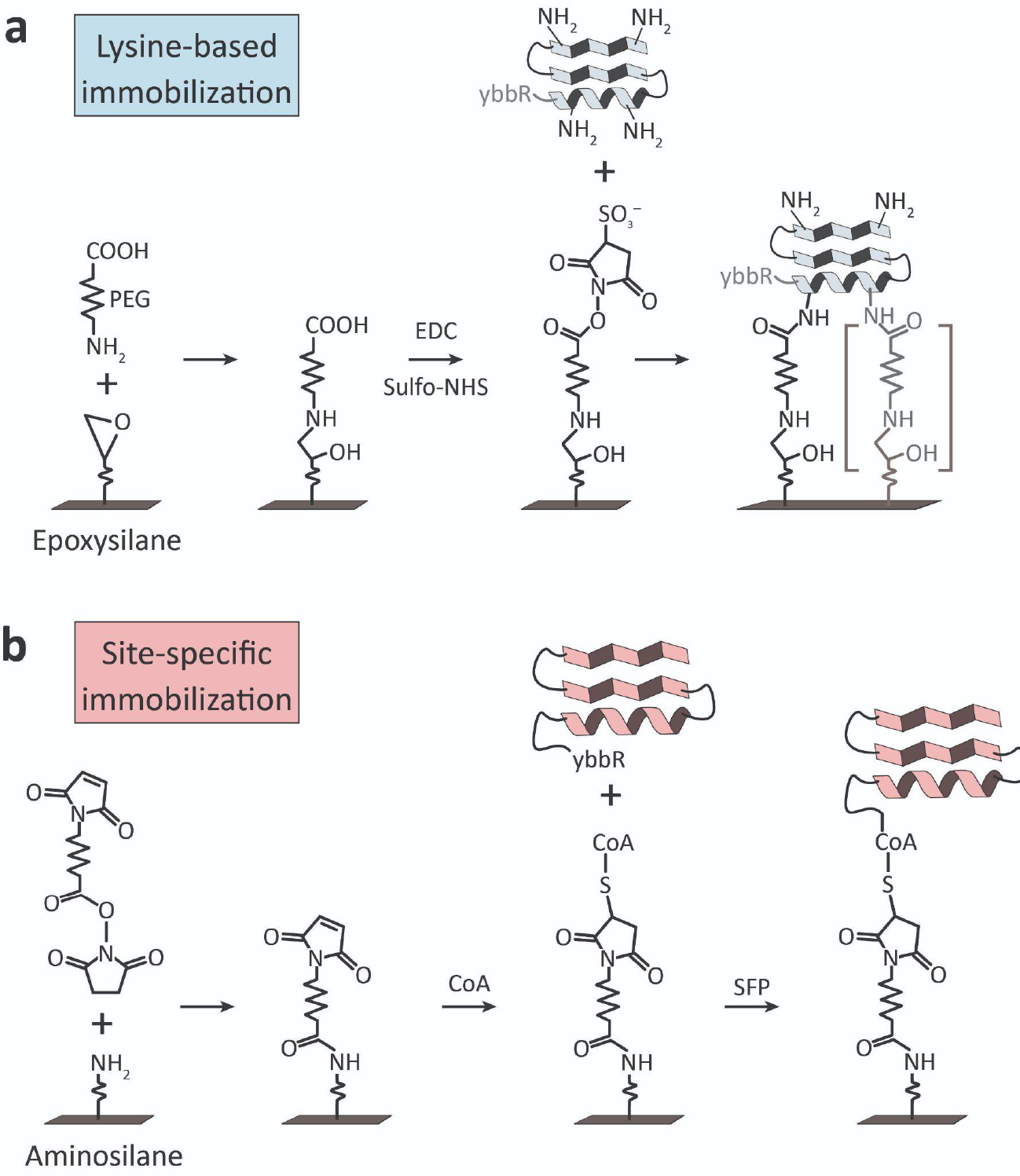
Surface chemistry used for lysine-based or site-specific protein immobilization in this study. (a) Protein immobilization through lysine groups was achieved by modifying epoxysilanized glass surfaces with heterobifunctional COOH-PEG-NH_2_. Following PEGylation, EDC/Sulfo-NHS was used to activate PEG carboxyls and couple to lysine groups on the protein of interest. Lysine-based immobilization resulted in a heterogeneity of tether points and shortening of the effective PEG linker length (right, grey). (b) Site-specific protein immobilization through a terminal ybbR tag was achieved by modifying aminosilanized glass surfaces with heterobifunctional maleimide-PEG-NHS. Following PEGylation, the sulfhydral group on coenzyme A (CoA) was coupled through reaction with maleimide. Exposure of the protein of interest bearing a terminal ybbR tag (DSLEFIASKLA) in the presence of the Sfp phosphopantetheinyl transferase site-specifically and covalently coupled the protein to the surface.

**Figure 2.**
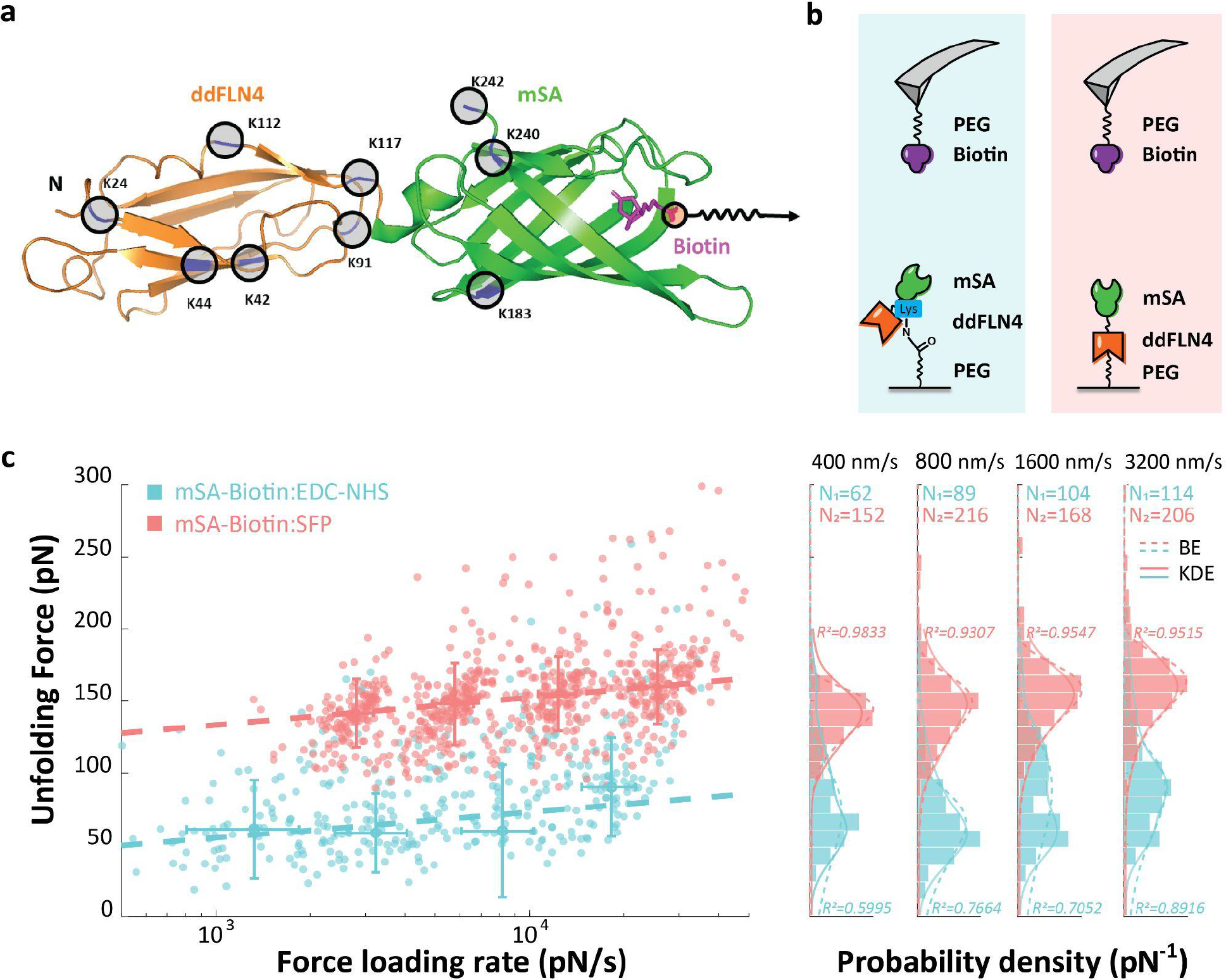
AFM-SMFS measurement on rupture force of mSA-Biotin complex using EDC/NHS or Sfp surface chemistry: (a) crystal structure (model) of ddFLN4-mSA construct showing surface lysine residues. (b) AFM setup showing the pulling geometry. (c) AFM-SMFS measurements showing rupture forces of the mSA-Biotin complex using lysine-based EDC/NHS or Sfp/ybbR tag surface chemistry at multiple pulling speeds. Dashed lines shown in the left panel are least square fits to the Bell-Evans model. Error bars represent the full width at half maximum (FWHM) of the probability distribution. The Bell-Evans energy landscape parameters from the fitting were Δx=0.52 ± 2.1 nm, ln(*k*_*o*_)= -2.04 ± 9.48 (R^2^=0.5386) for the EDC/NHS method and Δx=0.50 ± 0.09 nm, ln(*k*_*o*_)= -11.25 ± 3.61 (R^2^=0.9953) for the Sfp method. The histogram of rupture forces at different pulling speeds are shown in the right panel, fitted with kernel density estimates (KDE, shown in solid line) and Bell-Evans model prediction (dashed line with fitting R^2^ given). The most probable rupture forces from fitted KDE distribution under four pulling speeds with ± asymmetric full width at half maximum (FWHM) are shown in detail in Supplementary Table 1.

We probed mSA-biotin complex rupture events using AFM-SMFS at constant pulling speeds of 400, 800, 1600 and 3200 nm/s (**Fig. 2c**). The selection of valid force extension curves was performed under criteria of fitting force-extension behavior with a WLC model using a threshold force and distance value for the minimum adhesion force. In the case of site-specific mSA immobilization, an additional filter was applied based on fingerprint unfolding of ddFLN4 with characteristic contour length increment ∼32 nm from a two-step unfolding pattern, as a marker of single molecule interactions, so that non-specific adhesion was filtered out from the measurement. This type of filtering for fingerprint domain unfolding was not possible for the lysine-based method. For the lysine-based AFM measurement, we acquired 12,040 force-extension curves over a measurement time of 18 hours, resulting in 359 curves that passed our contour length filter (3.0%). For the ybbR site-specific immobilization, we acquired 13464 force-extension curves in 18 hours and obtained 742 usable curves, for a curve yield of 5.6%. This indicated that even though we used a more stringent filter based on fingerprint domain unfolding for the site-specific method, we obtained a higher number of high-quality usable curves. The rupture force histograms (**Fig. 2c, right**) demonstrate that the force distribution had higher magnitude and narrower width in the case of site-specific mSA immobilization as compared with lysine-based immobilization. The average full width at half maximum (FWHM) from fitted KDE distribution under four pulling speeds is 48.2pN for the Sfp method and 67.9pN for the EDC/NHS method. We also found that the site-specific method afforded rupture force distributions that more closely followed a theoretical two-state Bell-Evans model prediction (average fitting R^2^ under all pulling speeds for Sfp is 0.9551, compared to average R^2^=0.7399 for EDC/NHS method, shown in Fig.2.c right panel), with a left-skewed unimodal distribution, e.g. under a pulling speed of 400 nm/s, most probable rupture force ± asymmetric FWHM is: site-specific: 141.8 [+19.9 / -23.7] pN; lysine-based: 60.4 [+42.3 / -21.8] pN (**Sup. table 1**). The rupture forces measured using the Sfp method are found to be in a comparable range around 150 pN at force loading rates of 1E4 pN/s as for previously published under the same experiment configuration^[42]^.

### Resolving multiple dissociation pathway in SMFS

Next we compared lysine-based and site-specific immobilization for a protein-protein interaction that can unbind along different reaction pathways. We sought to understand how the coupling chemistry influences our ability to observe and classify different unfolding paths. We studied the multi-domain Xmodule-Dockerin/Cohesin (XDoc-Coh) receptor-ligand complex derived from the cellulolytic complex of *Ruminococcus champanellensis (Rc)* ^*[37]*^ (**Fig. 3**), which exhibits three distinct unfolding/unbinding pathways when dissociated under force, arising from the presence of dual binding modes in the complex and unfolding of the allosteric X-module domain. Following X-module unfolding, the binding interface is significantly destabilized, giving rise to a low force rupture event. Multi-pathway mechanical dissociation has been shown for a homologous XDoc/Coh system, and several distantly related Coh-Doc pairs were reported to have dual binding modes that give rise to multi-pathway dissociation reactions^[37,47,48]^.

**Figure 3.**
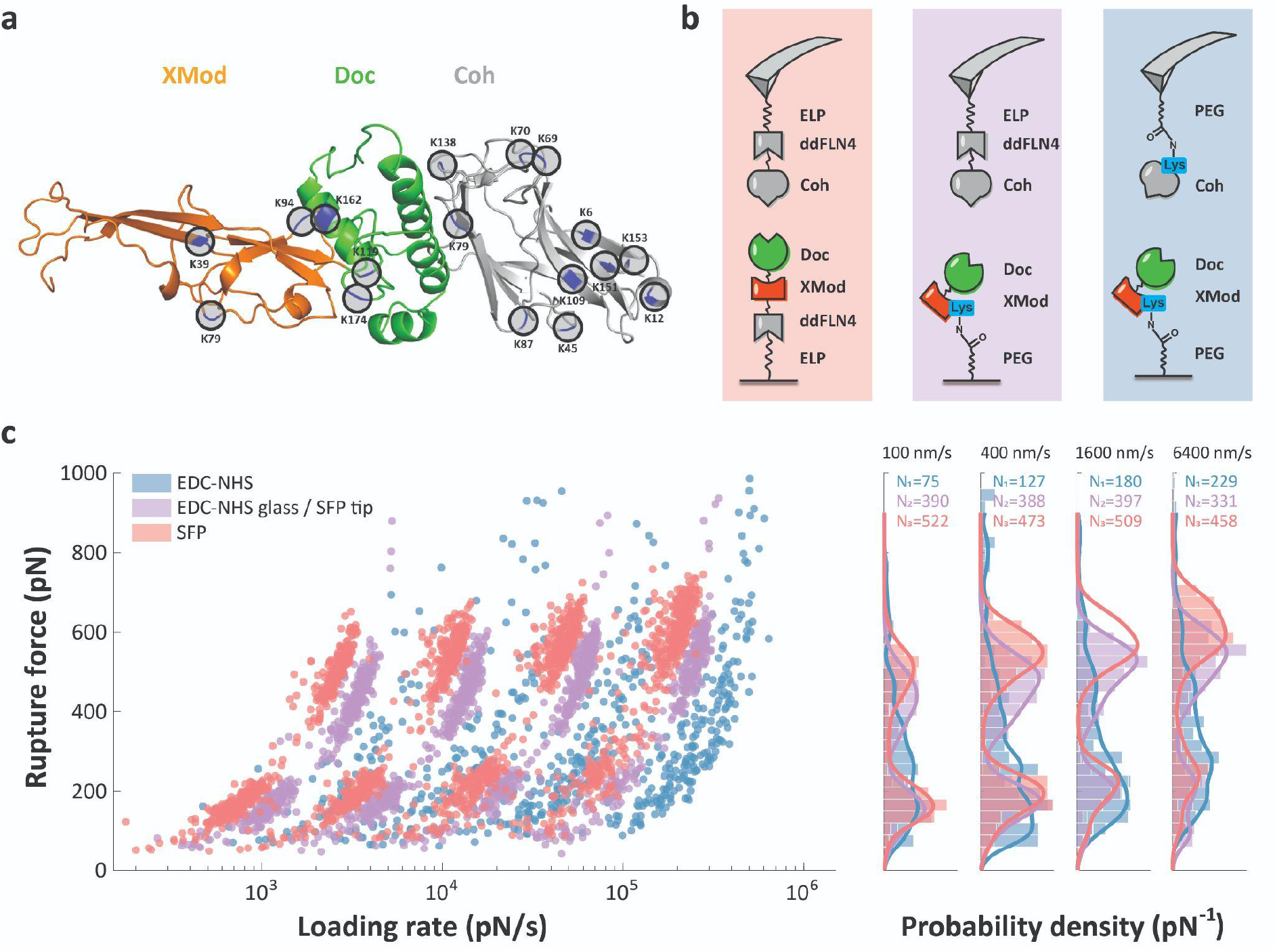
AFM-SMFS measurement on rupture force of XDoc-Coh complex using EDC/NHS or Sfp surface chemistry: (a) crystal structure of XDoc-Coh constructs showing potential binding sites. (b) AFM setup showing the pulling geometry. (c) Dynamic force spectra (left) and rupture force histograms (right) for the three measurement configurations show that site specific coupling best resolved the two rupture populations. The various pathways are shown in detail in Sup. Fig. 1.

The three signature unfolding pathways are shown in **Sup. Fig. 1**. In all three pathways, the final peak in the data trace represents breaking of the binding interface between Doc and Coh. We analyzed this final rupture event as a function of the loading rate and found substantial differences depending on the immobilization chemistry that was used. In this evaluation, we conducted AFM-SMFS on the XDoc-Coh complex using three coupling techniques: (i) site-specific immobilization on both cantilever and glass surface (**Fig. 3**, pink); (ii) lysine-based immobilization on both cantilever and glass surface (**Fig. 3**, blue); and (iii) a mixed method where ybbR/Sfp site-specific coupling was used to attach Coh to the cantilever and lysine-based coupling was used to attach XMod-Doc to the glass surface (**Fig. 3**, purple). For the Sfp site-specific coupling method with a fixed pulling terminus, the PEG linker was replaced by an ELP linker included in the fusion protein, which yields a better precision of the contour length for the pathway recognition and avoids PEG mechanical isomerization. These results show that when we used site-specific immobilization (**Fig. 3c**, pink), we could clearly resolve two rupture force populations (high-force pathway 1 and low force pathways 2 and 3), as well as clearly differentiate the two unfolding pathways among the low-force events where one low force pathway (pathway 2) included unfolding of XMod (**Sup. Fig. 1a**). When using lysine-based immobilization on both the cantilever and glass surface, we found that the high-force and low-force populations were not resolved and the distributions were broad and poorly defined (**Fig. 3c**, blue). When using the mixed surface chemistry approach, we obtained distributions with two observable discrete force populations, however the high force had a lower magnitude and higher loading rate attributed to the shortened effective linker length (see below).

At a given pulling speed, the lysine-based method resulted in higher loading rates (**Fig. 3c**). We also found that the complex rupture events occurred at shorter contour length for the lysine-based immobilization method (**Fig. 4**). The contour length at the time of rupture for the XMod-Doc/Coh system was found to be ∼20 nm, which is significantly shorter than the expectation for two times the PEG linker length (∼120 nm). This implies that multiple lysine residues were coupled to PEG chains, effectively shortening the linker length in the system.

**Figure 4.**
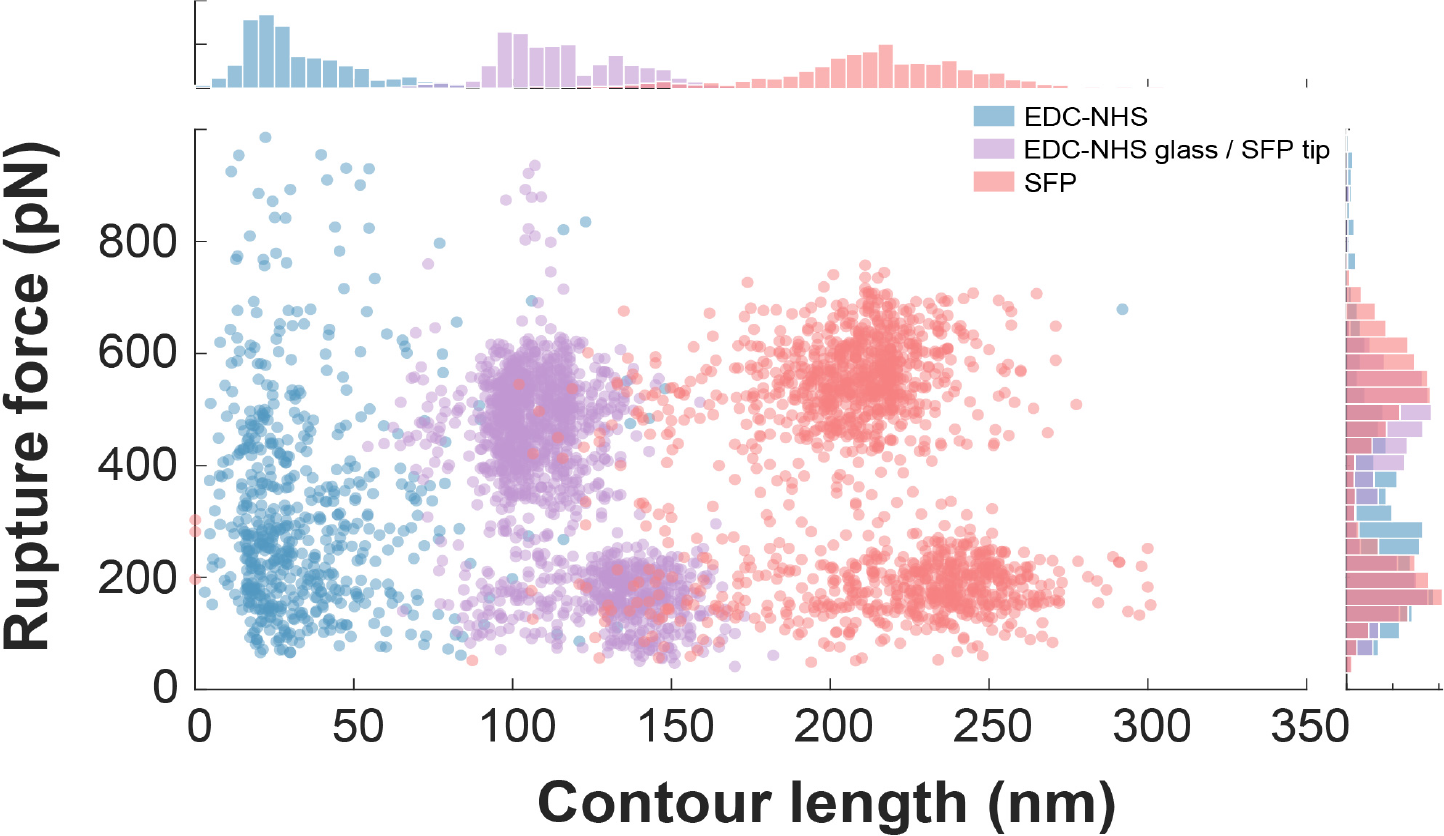
Comparison of contour length fitted from the AFM-SMFS measurement on rupture of XDoc-Coh complex using EDC/NHS or Sfp surface chemistry.

### Improved yield of useable data traces by site-specific coupling

Next we quantified the influence of site-specific methods in AFM-SMFS on yield and reproducibility of usable single-molecule traces. In AFM-SMFS, many of the acquired traces must be discarded because they do not represent the response of a single-molecule, and are difficult to interpret when multiple molecules are pulled in parallel. We found that the yield of single-molecule traces was significantly improved using the site-specific method. Under our measurement conditions using site-specific ybbR/Sfp coupling followed by an 18-hour AFM-SMFS measurement, around 20,000 approach-retract cycles on XMod-Doc/Coh were obtained. Among these, more than 2,000 (>10%) force-extension curves passed the analysis filter and represented single receptor-ligand rupture events. When the site-specific immobilization method was used, valid force curves could be precisely identified using the ddFLN4 domain as an unfolding fingerprint. For the lysine-based experiments, the selection of the force curves depends on ambiguous criteria including max adhesion force, distance of the rupture event and the force-extension behavior fitted using a WLC model. As a comparison, only around 200 curves were able to pass the selection filters for further analysis using the lysine-based EDC-NHS immobilization technique. This was due to a general low quality and short rupture distances of force-extension curves obtained using the lysine-based approach. The improved quality of the curves obtained with site-specific surface immobilization is further demonstrated in the rupture force vs. contour length plot (**Fig. 4**). This plot demonstrates that the final rupture event occurred farther away from the surface in the case of site-specific Sfp immobilization (>200 nm). When Xmodule-Dockerin/Cohesin complexes were probed using the mixed immobilization strategy (**Fig. 4, purple**), the contour length at rupture was at an intermediate length (100-150 nm). The slightly higher contour length at rupture for the low force rupture events in **Fig. 4** was due to the additional contour length increment upon XModule unfolding, which was characteristic of the low force pathway 2, and was observed for both site-specific immobilization and the mixed EDC-NHS/Sfp method. When only EDC/NHS was used on both the glass surface and the AFM cantilever tip, the contour lengths were severely shortened (<50 nm) even though the PEG linkers were of the same length. We attributed the shortened tether length to multiple tether points on the molecule (**Fig. 1a, right**).

### Compensation strategies to improve lysine-based methods

We attempted to improve the quality of the measurement for the lysine-based coupling method by minimizing the possibility of forming multiple PEG tethers that tended to shorten the system. We did so by diluting the surface density of the PEG linkers. The flexible linker included in the complex serves as an effective approach for the identification of molecular interactions at a single molecule level. Without flexible linker length, the measured complex rupture events are mixed with non-specific adhesion between the AFM tip and glass surface. We used a lower density of available binding sites by mixing the bifunctional COOH-PEG-NH_2_ active linker with a monofunctional methoxy-PEG-NH_2_ linker that lacks the carboxyl group required for EDC/Sulfo-NHS activation. As shown in **Sup. Fig. 4**, the contour length of the complex using partially non-functional methoxy-PEG-NH_2_ linkers was indeed higher, however, the curve yield in this case dropped even further to 142 curves that passed the analysis filter, obtained from 9,118 extend-retract cycles with the AFM (∼1.5%). The low yield was due to the high dilution of functional molecules on the surface, and the poor curve quality was not significantly improved in this case.

## Conclusions

Taken together, this direct comparison of lysine- vs. site-specific immobilization for mSA biotin and a multi-pathway XMod-Doc/Coh complex demonstrates the advantages of the site-specific approach in single-molecule mechanical assays. By anchoring the protein through a terminal ybbR tag, we were able to strictly control the molecular geometry of the tether point, avoid multiple tethers to the same molecule, and generally improve reliability and yield of the method. In cases where genetically encoded tags are not feasible (e.g., *in vivo*-derived samples), our results showed that a mixed site-specific/lysine-based method can be beneficial. Site-specific immobilization should therefore be used whenever possible in single-molecule force assays to improve precision, reliability, data quantity and reproducibility.

## Data Availability Statement

The data that support the findings of this study are openly available in Zenodo at [DOI], reference number [REF]. *** Correct DOI and reference number for dataset will be inserted prior to publication.

## Acknowledgements

This work was supported by the University of Basel, ETH Zurich, an ERC Starting Grant (MMA-715207), the NCCR in Molecular Systems Engineering, and the Swiss National Science Foundation (Project 200021_175478).

## Table of Contents

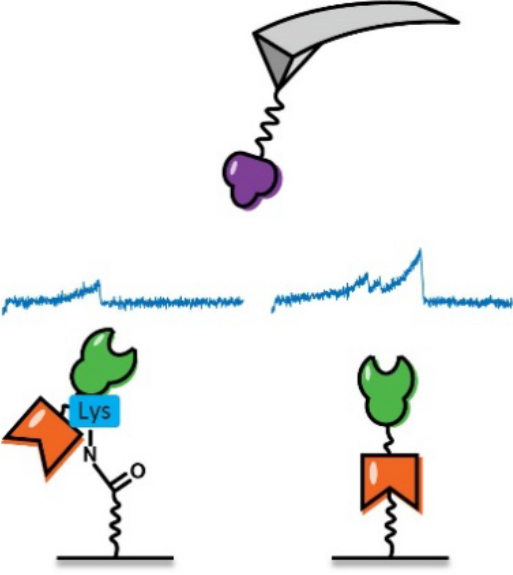

The surface immobilization technique for AFM-SMFS is crucial for both experimental yield, accuracy and resolution. In this paper we showed that a commonly used lysine-based immobilization can result in significant signal deterioration, including lowering of rupture forces, effective shortening of linkers, and loss of the ability to correctly classify unfolding pathways in a multi-pathway polyprotein system.

