## Supplementary Information for "Direct Comparison of Lysine vs. Site-specific Protein Surface Immobilization in Single-molecule Mechanical Assays"

[1] Department of Chemistry, University of Basel, 4058 Basel, Switzerland
[2] Department of Biosystems Science and Engineering, ETH Zurich, 4058 Basel, Switzerland
[3] Present address: Department of Bionanoscience, Delft University of Technology, 2629 Delft, The Netherlands
[4] National Center for Competence in Research, Molecular Systems Engineering, 4058 Basel, Switzerland
[5] Swiss Nanoscience Institute, 4056 Basel, Switzerland

**Abstract:** Single-molecule force spectroscopy (SMFS) is a powerful method for studying folding states and mechanical properties of proteins, however, it requires surface immobilization of proteins onto a force-transducing probe such as a cantilever or microscale bead. A common immobilization method relies on coupling surface-exposed lysine residues to carboxylated surfaces using 1-ethyl-3-(3-dimethyl-aminopropyl) carbodiimide and N-hydroxysuccinimide (EDC/NHS). Because proteins typically contain many lysine groups, this strategy results in a heterogeneous distribution of tether positions in the molecule. Genetically encoded peptide tags (e.g., ybbR) provide alternative chemistries for achieving site-specific immobilization, but thus far a direct comparison of site-specific vs. lysine-based immobilization strategies to assess effects on the observed mechanical properties was lacking. Here, we directly compared lysine- vs. ybbR-based protein immobilization in SMFS assays using several model polyprotein systems. Our results show that lysine-based immobilization results in significant signal deterioration for monomeric streptavidin-biotin interactions, and loss of the ability to correctly classify unfolding pathways in a multipathway Cohesin-Dockerin system. We developed a mixed immobilization approach where a site-specifically tethered ligand was used to probe surface-bound proteins immobilized through lysine groups, and found partial recovery of specific signals. The mixed immobilization approach represents a viable alternative for mechanical assays on in vivo-derived samples or other proteins of interest where genetically encoded tags are not feasible.

### Experimental Procedures

#### Protein construct designs, expression and purification

The constructs ybbr-HIS-ddFLN4-mSA, ybbr-ELP-ddFLN4-Xmod-Doc-HIS and Coh-ddFLN4-ELP-HIS-ybbr were designed for AFM measurements using the site-specific immobilization method. A ddFLN4 domain was inserted into a pET28a vector containing ybbr-HIS (for mSA), ybbr-HIS-ELP (for XMod-Doc) or ELP-HIS-ybbr (for Coh). Synthetic genes encoding mSA and XMod-Doc were inserted at the C-terminus of ddFLN4 using Gibson assembly, respectively, and the Coh synthetic gene was inserted to the N-terminus of ddFLN4 using restriction digestion cloning (NdeI and BamHI sites). For AFM measurements on XDoc-Coh using the lysine-based immobilization method, ELP and ddFLN4 domains were removed from the construct. These sequences were confirmed by Sanger sequencing (Microsynth AG).

The protein samples used for AFM measurement were expressed using NiCo21(DE3) competent cells (New England Biolabs, Ipswich, MA, USA). The cells were grown in terrific broth (TB) containing 50 µg/mL kanamycin at 37 °C until OD reached ~ 0.6. The protein expression was induced by 0.5 mM of isopropyl β-D-1-thiogalactopyranoside (IPTG) and incubated at 20 °C overnight with shaking. The cells were subsequently pelleted and lysed using sonication. The cell lysate was loaded to a His-trap column (GE Healthcare, IL, USA), washed with phosphate buffered saline (PBS, 137 mM NaCl, 2.7 mM KCl, 10 mM $Na_2HPO_4$ and 2 mM $KH_2PO_4$, pH 7.4) with 20 mM imidazole and eluted with PBS buffer supplemented with 500 mM imidazole. The eluate was further purified with Superdex Increase 200 10/300 GL size-exclusion column (GE Healthcare).

#### Immobilization of ybbr-tagged protein for AFM-SMFS

Ybbr-tagged proteins were immobilized on functionalized AFM cantilevers and cover glasses according to previously published protocols[1,2]. In brief, coverglasses were first cleaned by UV-ozone treatment for 40 minutes followed by soaking in piranha etching solution and rinsing in water. Cantilevers were treated with UV-ozone. Next, levers and coverglasses were treated with 3-Aminopropyl (diethoxy) methylsilane (APDMES, ABCR GmbH, Karlsruhe, Germany) to silanize the surface with amine groups. The amine groups were subsequently conjugated to a heterobifunctional NHS-PEG-Mal linker (5 kDa, Rapp Polymere, Tübingen, Germany) in HEPES buffer (50 mM, pH 7.5) for 30 min, followed by incubating with Coenzyme A (CoA, 200 µM) in coupling buffer (50mM sodium phosphate, 50mM NaCl, 10mM EDTA, pH 7.2) for 2 hours at room temperature. For experiment with ELP linker included in the ybbr-tagged fusion protein, instead a small-molecule crosslinker (sulfosuccinimidyl 4-(N-maleimidomethyl)cyclohexane-1-carboxylate), sulfoSMCC), which added negligible contour length (0.83 nm) to the system was immobilized onto amino silanized surfaces. Finally, the ybbr-tagged proteins were covalently immobilized to the CoA surfaces or cantilevers by a Sfp-catalyzed coupling reaction in the measurement buffer (TBS, $Ca^{2+}$ for Coh-Doc, and PBS for mSA-Biotin) at room temperature for 2 hours. For the Biotin-functionalized cantilever, silanized cantilevers were incubated with 25 mM NHS-PEG-Biotin (5 kDa, Rapp Polymere, Tübingen, Germany) in 100 mM HEPES buffer for 30 min. Functionalized cantilevers and coverglasses were kept in the measurement buffer until the measurement.

#### Protein immobilization using EDC/NHS for AFM-SMFS

The epoxy surface was treated with 25 mM COOH-PEG-$NH_2$ (5 kDa, Rapp Polymere, Tübingen, Germany) in HEPES buffer (100 mM, pH 8.0) for 30 mins at room temperature. The NHS group was then activated by incubation with 40 mM EDC and 90 mM Sulfo-NHS in MES buffer (100 mM, pH 6.0) for 20 mins at room temperature. The protein solution was added and incubated for 2 hours in a wet chamber at room temperature. To quench the remaining NHS ester groups, the treated AFM tip and glass were rinsed in Tris buffer (TBS; 50 mM Tris, 150 mM NaCl, pH 7.6) for 20 min at room temperature.

#### AFM-SMFS measurement and data analysis

Force spectroscopy measurements were performed on a Force Robot AFM (JPK instruments, Berlin, Germany). Cantilever spring constants (ranging from 0.07 to 0.1 $N \cdot m^{-1}$) were calibrated using the contact-free method. The cantilever was brought into contact with the surface and withdrawn at constant speed ranging from 100 to 6400 $nm \cdot s^{-1}$. During the measurement, the force curves collected were first analyzed and filtered in a real-time manner by software available on the AFM instrument with a loose criteria to exclude plain force traces and nonspecific binding to the glass substrate: the max adhesion force of retraction segment with a minimum extension of 10 nm should be larger than 30 pN and number of force peaks from WLC chain model fitting is between 1 and 10. After the measurement, the selected force curves showing positive force signals were processed and analyzed by contour length transformation[2]. For Sfp surface chemistry with a fingerprint domain of ddFLN4 included, the single molecular force traces were identified by searching for contour length increments that matched the fingerprint domain of ddFLN4 (≈36 nm). For EDC/NHS methods where fingerprint domains are not available, the force traces were examined by checking the number of force peaks from the contour length histogram that represent the complex rupture and the possible unfolding of the protein domain included in the construct.

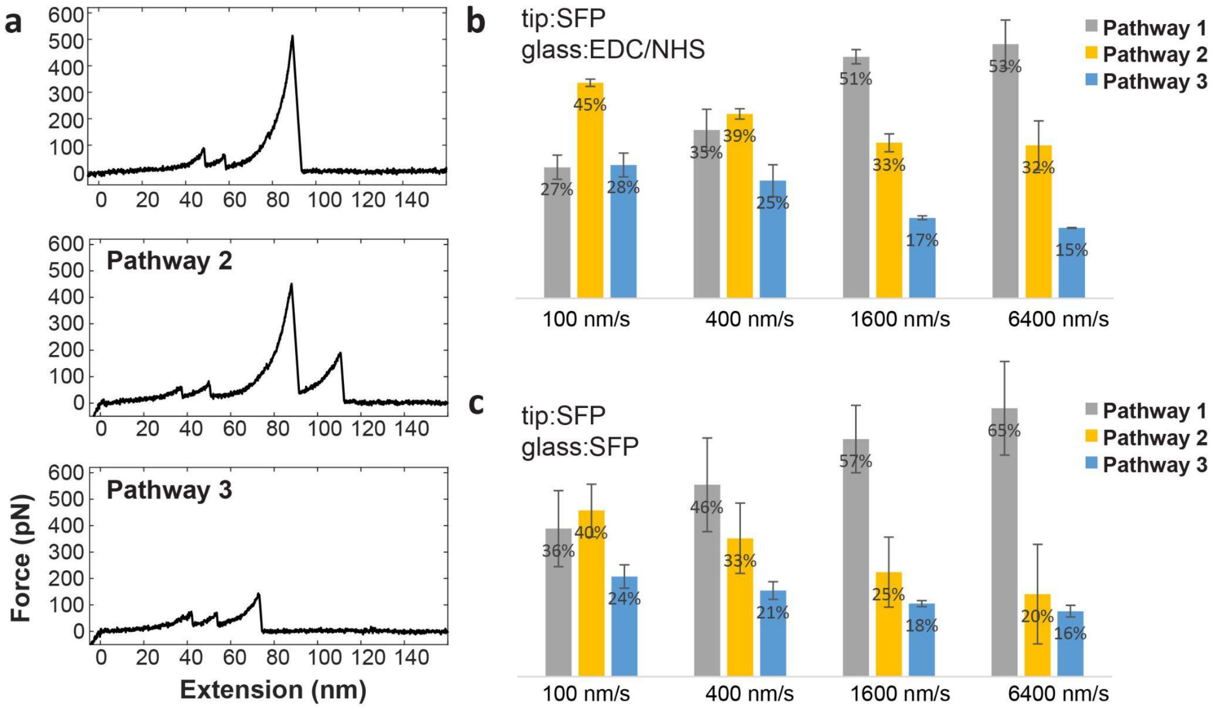


**Supplementary Figure 1.** Pathway ratios obtained from AFM-SMFS measurements on XMod-Doc/Coh complex using EDC/NHS or Sfp surface chemistry: (a) Representative force curves showing three pathways: Pathway 1, high force; Pathway 2, Xmod unfolding; Pathway 3, low force. Distribution of different pathways are obtained from AFM-SMFS measurements using (b) Sfp surface chemistry for AFM tip and EDC/NHS method for the glass surface and (c) Sfp for both sides. In (b) and (c) the average pathway ratio from 2 sets of measurement are given and the error bars represent the standard deviation for each group. When EDC/NHS was used on both the cantilever tip and coverglass, it was not possible to classify pathways.

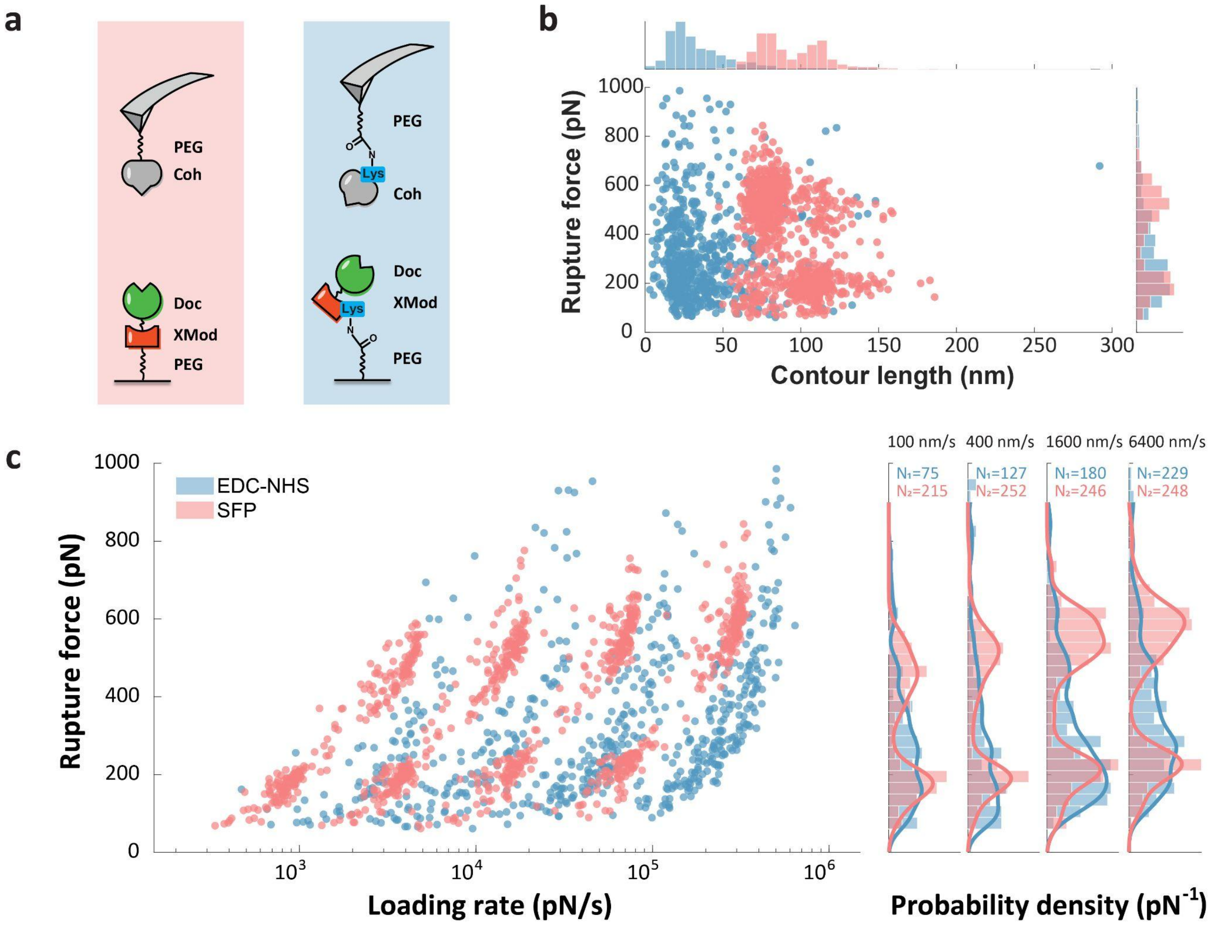


**Supplementary Figure 2 .** AFM-SMFS measurement on rupture force of XDoc-Coh complex using EDC/NHS or Sfp surface chemistry: (a) AFM setup showing the pulling geometry: identical structures including the same PEG linkers are used. (b) Fitted of contour length of XDoc-Coh complex rupture events. (c) Dynamic force spectra (left) and rupture force histograms (right) for the three measurement configurations show that site specific coupling best resolved the two rupture populations.

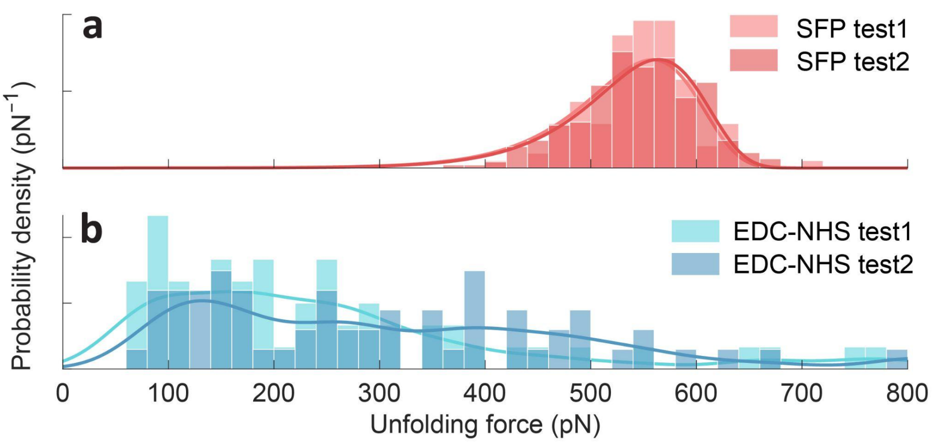


**Supplementary Figure 3.** Reproducibility of AFM-SMFS measurement on rupture force of XDoc-Coh complex using (a) Sfp and (b) EDC/NHS surface chemistry. AFM data sets were obtained on two pairs of samples from overnight measurements under the same condition, with ~4000 total force traces collected within 18 hours for each sample. In a) Sfp group, the histogram is obtained from the high force rupture pathway, then fitted with BE model (solid line). The most probable rupture force is 558.3 [+51.4 -76.2] pN for replicate 1 (104 curves in total) and 563.9 [+51.2 -76.1] pN for replicate 2 (251 curves in total). In b) EDC-NHS group, the rupture forces could not be distinguished between high force and low force pathways. The rupture force histogram was then fitted with KDE and the most probable rupture force is 162.7 [+167.2 -116.9] pN for replicate 1 (60 curves in total) and 132 [+341.1 -65.1] pN for replicate 2 (67 curves in total).

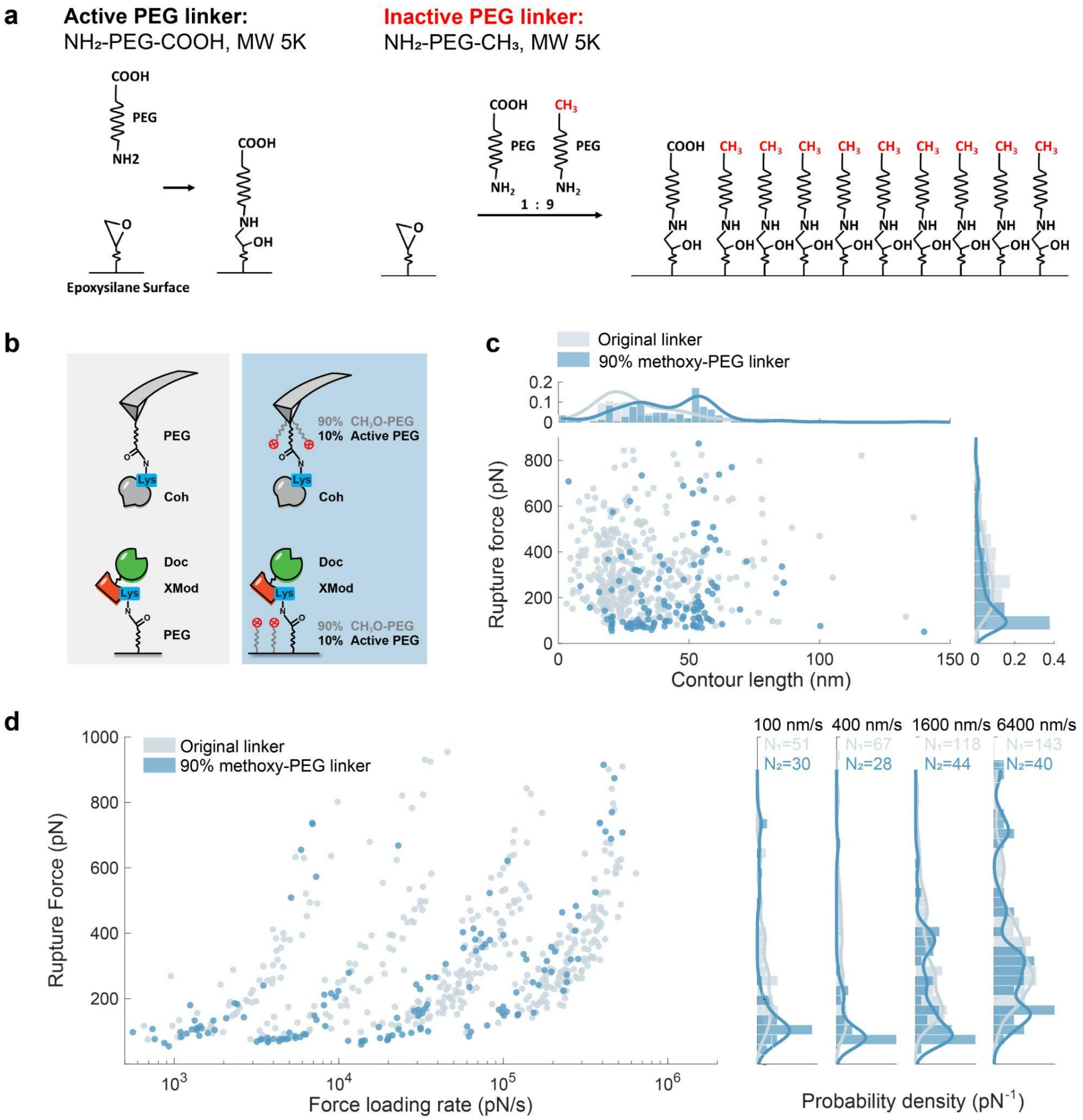


**Supplementary Figure 4.** AFM-SMFS measurements of XMod-Doc/Coh complex using EDC/NHS surface chemistry with PEG linker. (a) Schematic showing dilution of heterobifunctional PEG linker with monofunctional methoxy-PEG-$NH_2$ which can be bound to the surface but cannot be coupled to the protein. (b) Schematic of the AFM measurement setup comparing 100% functional PEG to 10% functional PEG on both the surface and cantilever. (c) Rupture force vs. contour length plot showing 100% vs. 10% functional linker. (d) Rupture force vs. log (loading rate) comparing 100% functional PEG to 10% functional PEG linker. For all plots, grey: 100% functional linker (COOH-PEG-$NH_2$); blue: 10% functional linker / 90 % non-functional (methoxy-PEG-$NH_2$).

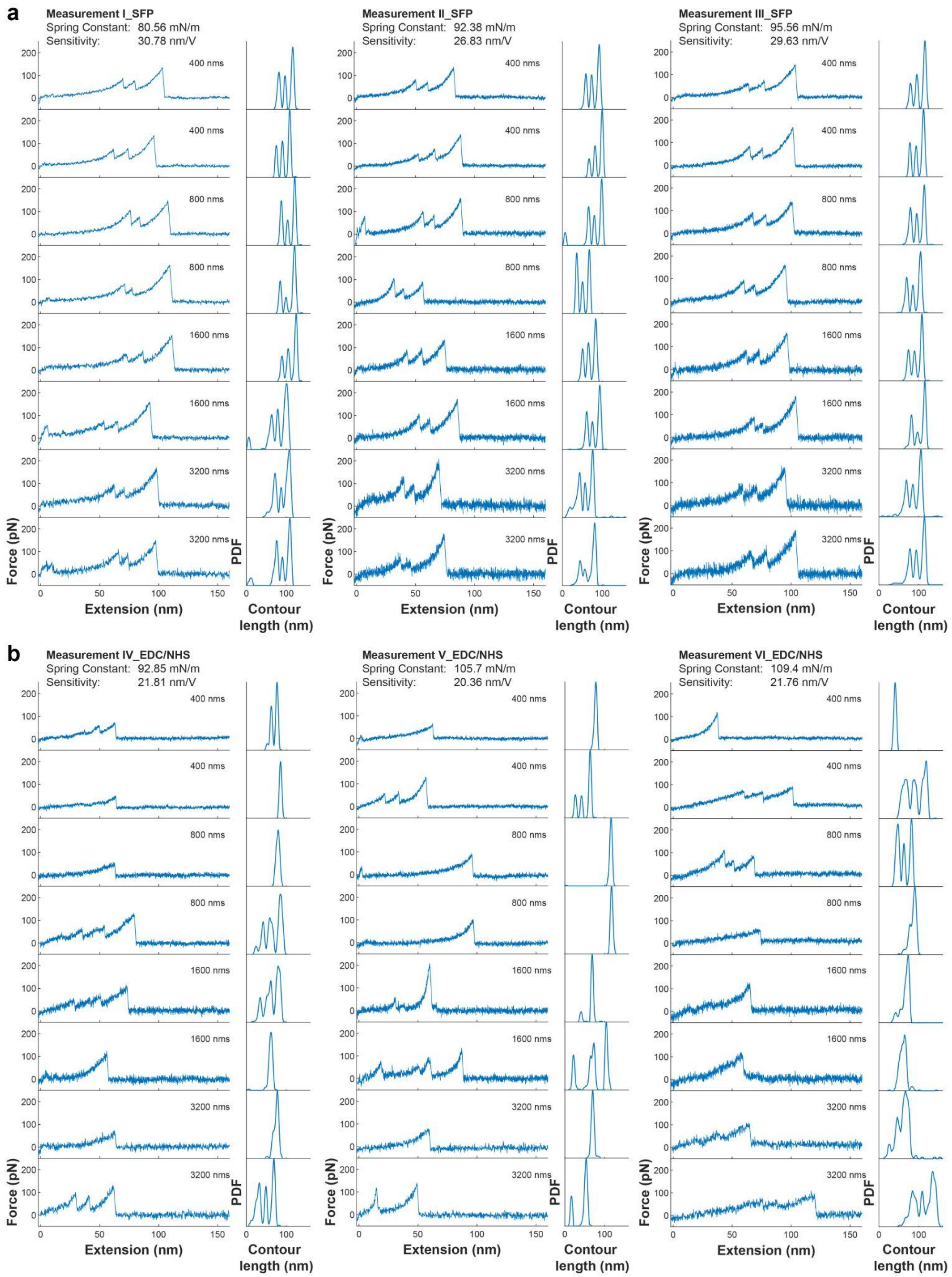


**Supplementary Figure 5.** Examples of raw AFM-SMFS force extension traces and corresponding contour length of mSA-Biotin complex pulling test under different measurement configuration: a) Sfp surface chemistry and b) EDC/NHS surface chemistry. For each group 3 replicate measurements using different samples are shown with cantilever information listed above.

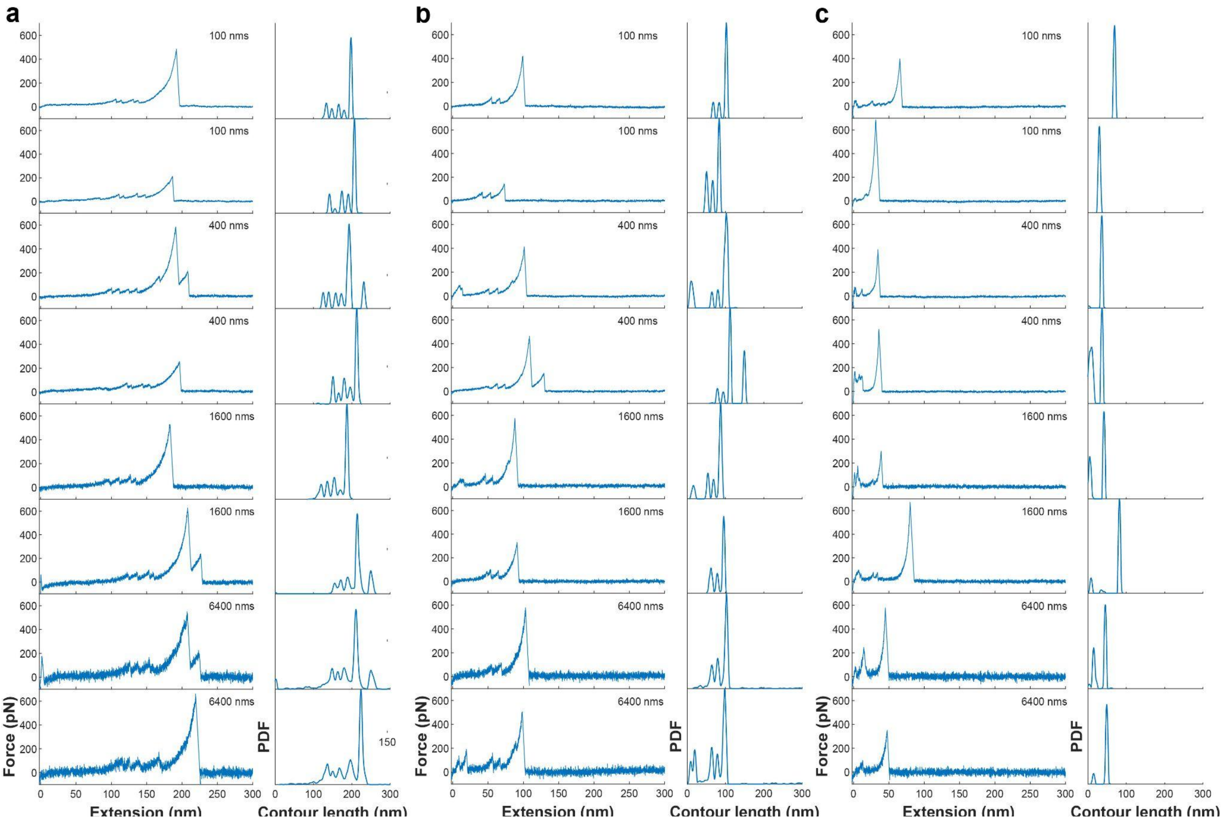


**Supplementary Figure 6.** Examples of AFM-SMFS force extension traces and corresponding contour length of Coh-Doc complex pulling test under different measurement configuration: **a)** Sfp surface chemistry, **b)** EDC/NHS surface and Sfp tip, and **c)** EDC/NHS surface chemistry

| | 400 nm/s | 800 nm/s | 1600 nm/s | 3200 nm/s |
|---|---|---|---|---|
| **Sfp** | 141.8 pN<br>[+19.9 / -23.7] pN | 148.3 pN<br>[+22.3 / -30.8] pN | 155.6 pN<br>[+21.2 / -26.8] pN | 159.6 pN<br>[+23.9 / -24.1] pN |
| **EDC/NHS** | 60.4 pN<br>[+42.3 / -21.8] pN | 57.8 pN<br>[+27.9 / -23.3] pN | 58.6 pN<br>[+69.6 / -20.6] pN | 90.9 pN<br>[+23.3 / -42.5] pN |

**Table S1.** The most probable rupture force of rupture force of mSA-Biotin complex (shown in Figure 2) from fitted KDE distribution under four pulling speeds with ± asymmetric full width at half maximum (FWHM) for the Sfp method and for the EDC/NHS method.

| dx | ln(k0) | k0 | Ref. |
|---|---|---|---|
| 0.50 nm | -11.25 | 1.30E-05 $s^{-1}$ | Fig1. |
| 0.41 nm | -8.14 | 2.92E-04 $s^{-1}$ | Santos et al. 2022[3] |
| 0.39 nm | -11.33 | 1.20E-05 $s^{-1}$ | Bauer et al. 2018[4] |

**Table S2.** Comparison of fitted parameters of complex rupture on an identical construct of mSA-Biotin with N-terminal pulling configuration using AFM-SMFS in previous studies. The obtained rupture force and loading rate under different pulling speeds were fitted with Bell-Evans model and the barrier distance dx as well as the off-rate k0 were yielded form the fitting process.

**Protein sequences for AFM-SMFS including marker domains**

**ybbr-HIS- ddFLN4- mSA**

**DSLEFIASKLA**HHHHHH**GS**ADPEKSYAEGPGLDGGECFQPSKFKIHAVDPDGVHRTDGGDGFVVTIEGPAPVDPVM
VDNGDGTYDVEFEPKEAGDYVINLTLDGDNVNGFPKTVTVKPAP**GSGSGSGS**AEAGITGTWYNQSGSTFTVTAGAD
GNLTGQYENRAQGTGCQNSPYTLTGRYNGTKLEWRVEWNNSTENCHSRTEWRGQYQGGAEARINTQWNLTYEG
GSGPATEQGQDTFTKVK

**Coh-HIS-ybbr**

**ADGAAKLSMDQKFAEPGETVEIALNLENFDASWTGLEFLVNYDPKLEVALDGAGDIDYSYGDAIGAMGKKISVGG
AISKDLTADGLKGFAFAWGTATAISGNGQLGVFKFTVPADAQPGDEFPVNLTVNVGSFIDANKENIPFETVNGWI
KIKEEGS**HHHHHH**GS**DSLEFIASKLA

**Coh-ddFLN4-ELP-HIS-ybbr**

**ADGAAKLSMDQKFAEPGETVEIALNLENFDASWTGLEFLVNYDPKLEVALDGAGDIDYSYGDAIGAMGKKISVGG
AISKDLTADGLKGFAFAWGTATAISGNGQLGVFKFTVPADAQPGDEFPVNLTVNVGSFIDANKENIPFETVNGWI
KIKEEGSGSGSGS**ADPEKSYAEGPGLDGGECFQPSKFKIHAVDPDGVHRTDGGDGFVVTIEGPAPVDPVMVDNGD
GTYDVEFEPKEAGDYVINLTLDGDNVNGFPKTVTVKPAP**GSGSGSHG**VGVPGMGVPGVGVPGVGVPGVGVPGVG
VPGVGVPGVGVPGVGVPGEGVPGEGVPGVGVPGMGVPGVGVPGVGVPGVGVPGVGVPGVGVPGVGVPGVGV
PGEGVPGEGVPGVGVPGMGVPGVGVPGVGVPGVGVPGVGVPGVGVPGVGVPGVGVPGVGVPGEGVPGEGVPG**WRG**H
HHHHH**GS**DSLEFIASKLA

**ybbr-XMod-Doc-HIS**

**GT**DSLEFIASKLA**GSSGSETTTETSVSQSFSFTEPTRVCYWSHDNRPFDLEGLKVILQETIFIIHEDGTVVEAEEQPAAV
DITEQCMTIFCNDKQVATPEDTYETQIYKYMLEFAYEGNPDVIVGTHQVLIGVKGDTNLSGDVTVADAVEILMYR
ADMMANPNNPTYVFNEDPDLHKLGYFLGNVDCIKEGDDLNVADAVNILMFRAEVLANGSATWEQIVGYDLIDP
SDESTFRTIEKLGS**HHHHHH

**ybbr-ELP-ddFLN4-XMod-Doc-HIS**

**GT**DSLEFIASKLA**WGSGHG**VGVPGMGVPGVGVPGVGVPGVGVPGVGVPGVGVPGVGVPGVGVPGEGVPGEGV
PGVGVPGMGVPGVGVPGVGVPGVGVPGVGVPGVGVPGVGVPGVGVPGEGVPGEGVPGVGVPGMGVPGVGVP
GVGVPGVGVPGVGVPGVGVPGVGVPGVGVPGEGVPGEGVPG**WPSGS**ADPEKSYAEGPGLDGGECFQPSKFKIHA
VDPDGVHRTDGGDGFVVTIEGPAPVDPVMVDNGDGTYDVEFEPKEAGDYVINLTLDGDNVNGFPKTVTVKPAP**GS
GSGSGSSGSETTTETSVSQSFSFTEPTRVCYWSHDNRPFDLEGLKVILQETIFIIHEDGTVVEAEEQPAAVDITEQCM
TIFCNDKQVATPEDTYETQIYKYMLEFAYEGNPDVIVGTHQVLIGVKGDTNLSGDVTVADAVEILMYRADMMAN
PNNPTYVFNEDPDLHKLGYFLGNVDCIKEGDDLNVADAVNILMFRAEVLANGSATWEQIVGYDLIDPSDESTFRTI
EKLGS**HHHHHH
